# LieOTDock: A Protein Docking Framework via Lie Group Optimization and Optimal Transport

**DOI:** 10.1101/2025.09.04.674197

**Authors:** Yue Hu, Zanxia Cao, Yingchao Liu

## Abstract

Protein-protein interactions (PPIs) are fundamental to cellular function, and structurally characterizing them is a key goal in molecular biology. Computational docking, a crucial tool for predicting the structure of protein complexes, faces significant challenges in search efficiency and scoring accuracy. Here, we introduce LieOTDock, a novel framework that leverages the mathematics of Lie groups and Optimal Transport (OT) for highly efficient protein docking pose generation. The core of our method is a data-driven surface sampling technique that identifies salient geometric features for initial pairing. These initial poses are then refined via a gradient-based optimization directly on the Lie group SE(3), the natural space of rigid-body motions. This optimization is guided by a differentiable score based on Sinkhorn-regularized Optimal Transport, which measures the geometric complementarity of surface patches. We tested LieOTDock on the 1AVW benchmark, where it successfully generated a near-native pose of 3.43 Å RMSD from a 5000-candidate run in under three minutes on a consumer-grade Apple M4 Mini. Our results demonstrate that LieOTDock, by its principled use of Lie theory and OT, provides a powerful and efficient framework for generating high-quality candidate structures. This work decouples the challenge of geometric sampling from final scoring, offering a robust and modular approach to the protein docking problem, though future work is needed to integrate more physically realistic scoring functions for improved candidate ranking.

## 1 Introduction

Protein-protein interactions (PPIs) are fundamental to nearly all cellular processes, from signal transduction and metabolic regulation to immune responses. Understanding the three-dimensional structure of protein complexes is therefore a cornerstone of modern molecular biology, providing critical insights into biological function and paving the way for rational drug design [1]. While experimental methods like X-ray crystallography and cryo-electron microscopy provide high-resolution structural data, they are often time-consuming and technically challenging. Consequently, computational protein-protein docking has emerged as an indispensable tool for predicting the structure of these complexes [2, 3].

The protein docking problem is notoriously difficult, primarily due to two major challenges: the vastness of the six-dimensional translational and rotational search space, and the complexity of accurately scoring and ranking the generated candidate poses [4]. Mainstream approaches often rely on grid-based methods using Fast Fourier Transforms (FFT) to rapidly evaluate shape complementarity, as exemplified by programs like ZDOCK [5]. While successful, these methods can be computationally intensive and sensitive to the fine details of surface topology. The search for a correct binding pose is a global optimization problem, but the subsequent evaluation and ranking of these poses require a scoring function that can accurately approximate the binding free energy, a task that remains a major bottleneck in the field [6].

To address the search efficiency challenge, we developed LieOTDock, a novel framework for protein docking that prioritizes computational efficiency and robust geometric sampling through the application of modern mathematical tools. The core contribution of our work is the synergy of three key ideas. First, we use a novel, data-driven method for sampling a molecular surface to identify salient features for docking, which are scored continuously based on their local geometry. Second, by pairing these complementary features, we generate a diverse set of high-quality initial poses. Third, and most critically, each pose is then refined using a gradient-based optimization directly on the Special Euclidean Lie group SE(3), the natural manifold of rigid-body motions. This optimization is guided by a local alignment score derived from the principles of Optimal Transport (OT), specifically using the computationally efficient Sinkhorn algorithm [7]. This Lie-theoretic approach allows for unconstrained optimization in the tangent space of the group, avoiding singularities associated with other representations of rotation.

This paper details the LieOTDock framework, from surface generation to final pose refinement. We demonstrate that LieOTDock is a highly effective pose generator, capable of producing near-native structures that can then be evaluated by external, more sophisticated scoring functions. Our work effectively decouples the difficult problem of global scoring from the initial geometric search, offering a modular and extensible platform for future development in protein docking.

## 2 Methods

Our docking framework, LieOTDock, is a multi-stage process designed to efficiently sample a wide range of orientations and generate high-quality candidate poses. The workflow is divided into three main phases: (1) molecular surface generation and feature identification, (2) pose generation and refinement using Lie group optimization guided by an Optimal Transport score, and (3) final evaluation against a reference structure.

### 2.1 Molecular Surface and Feature Point Generation

The initial step involves translating the three-dimensional atomic coordinates of the receptor and ligand from their PDB files into a continuous and differentiable representation of the molecular surface.

#### 2.1.1 PDB Parsing

The first step in our pipeline is parsing the atomic coordinates from PDB files for both the ligand and receptor molecules. For each protein, we extract the coordinates of all heavy atoms, their element types to assign van der Waals (VdW) radii, and their residue and atom names to assign partial charges. A key feature of our parser is the filtering of alternate locations (we only consider atoms with altLoc ‘ ‘ or ‘A’) and the exclusion of hydrogen atoms to simplify the model. For RMSD calculations, we also extract the coordinates of the C-alpha atoms to form a backbone representation.

#### 2.1.2 Surface Triangulation

We employ the MSMS (Michel Sanner’s Molecular Surface) program [8] to generate a solvent-excluded surface (SES) point cloud. Given the atomic coordinates and their respective van der Waals (VdW) radii, MSMS computes a triangulated mesh representing the protein’s surface. This process yields a set of *n* surface vertices𝒱 = {**v**_1_, …, **v**_*n*_ } ⊂ ℝ^3^and their corresponding outward-facing normal vectors 𝒩= {**n**_1_, …, **n**_*n*_ } ⊂ ℝ^3^, where each ∥ **n**_*i*_ ∥= 1. The density of the vertex mesh is controlled by the `‘msms density’` parameter.

#### 2.1.3 Geometric Feature Scoring

Instead of relying on discrete classifications of surface shape, we implemented a continuous scoring function to identify salient geometric features, inspired by the principles of local curvature and solvent exposure. This method assigns a unique feature score *S*_*f*_ to each vertex **v**_*i*_ on the molecular surface, allowing for a robust ranking of all points from most concave (pit-like) to most convex (peak-like).

For each vertex **v**_*i*_ ∈ 𝒱, the algorithm proceeds as follows:

1. **Neighbor Identification:** We first identify the set 𝒦 _*i*_ of the *k* nearest neighboring vertices to **v**_*i*_ using a *k*-d tree for efficient querying. The number of neighbors *k* is a tunable parameter.
2. **Depth Score Calculation:** A “depth score” (*S*_depth_) is computed, which measures the average displacement of neighboring points along the normal vector **n**_*i*_. Let **d**_*ij*_ = **v**_*j*_ − **v**_*i*_ be the vector from the central point **v**_*i*_ to a neighbor **v**_*j*_ ∈ 𝒦_*i*_. The depth score is the mean of the projections of these vectors onto the normal:

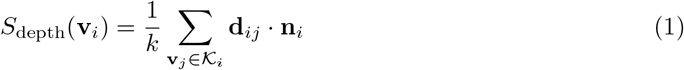 A negative *S*_depth_ indicates that the neighbors are, on average, located “in front of” the vertex along its normal, characteristic of a concavity or pit. Conversely, a positive score indicates a convexity or peak.
3. **Wideness Score Calculation:** To complement the depth, a “wideness score” (*S*_wideness_) is calculated as the sum of the standard deviations of the neighbor coordinates. This measures the spatial dispersion of the local patch, providing a sense of its breadth.
4. **Final Feature Score:** The final feature score *S*_*f*_ for the vertex **v**_*i*_ is the product of the depth and wideness scores:

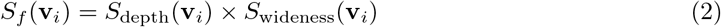

This combined score prioritizes features that are both deep/high and broad, which are often key to biomolecular recognition.

After computing *S*_*f*_ for all vertices, we sort the vertices based on this score. The *N*_feat_ vertices with the lowest scores are designated as the set of concave features, ℱ_*c*_, and the *N*_feat_ vertices with the highest scores are designated as the set of convex features, ℱ_*v*_.

### 2.2 Pose Generation and Refinement

With salient features identified on both the receptor and ligand surfaces, we generate a large set of candidate docking poses by aligning complementary patches. The refinement of these poses is where the core mathematical framework of our method is applied.

#### 2.2.1 Pose Refinement on the Lie Group SE(3)

The position and orientation of a rigid body, such as a protein, can be described as an element of the Special Euclidean group, SE(3). This is a Lie group that combines 3D rotations (from the Special Orthogonal group SO(3)) and 3D translations. A key insight from robotics and computer vision is that performing optimization directly on this curved manifold is more robust than using parameterizations like Euler angles or quaternions, which can suffer from singularities or constraints [9].

Instead of optimizing the 4×4 transformation matrix directly, we optimize in the associated Lie algebra, 𝔰 𝔢 (3). The Lie algebra is a 6-dimensional vector space, and its elements, known as “twists” or “screws”, represent infinitesimal rigid motions (3 for rotation, 3 for translation). We can map an element ***ξ*** ∈ 𝔰 𝔢 (3) from the algebra to a transformation **T** ∈ *SE*(3) using the matrix exponential map, **T** = exp(***ξ***) [10]. This allows us to use standard unconstrained gradient-based optimization methods.

For each initial pose, we perform a local refinement to maximize the surface complementarity. This is framed as an optimization problem to find a local rigid transformation **T**_*align*_ ∈ *SE*(3) that maximizes a score. The transformation is parameterized by a 6-dimensional twist vector ***ξ*** ∈ 𝔰 𝔢 (3), and the optimization is performed using the AdamW optimizer.

#### 2.2.2 Local Alignment Score via Optimal Transport

Each initial pose is subjected to a local refinement to optimize the complementarity of the local surface patches. A patch is defined as all surface vertices within a given textttpatch radius of a feature point.

This refinement is guided by a Sinkhorn alignment score, which is derived from the principles of regularized Optimal Transport [7]. The score *S* between a receptor patch 𝒫_*r*_ and a transformed ligand patch 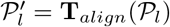 is defined as:

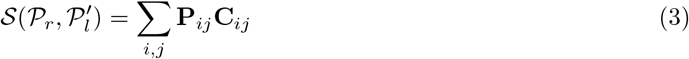

where **C** is a cost matrix encoding rewards for proximity and penalties for steric clashes:

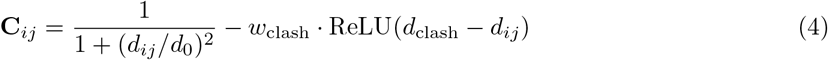

Here, *d*_*ij*_ is the Euclidean distance between points **v**_*i*_ ∈ 𝒫_*r*_ and 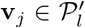 The matrix **P** is the result of applying a set number of Sinkhorn normalization iterations to exp(*γ***C**), which yields a soft assignment, or “transport plan”, between the points of the two patches. This score is fully differentiable with respect to the ligand’s pose, allowing for efficient gradient-based optimization.

The final transformation for each trial is the composition of the initial and alignment transformations: **T***_3_*= **T***_3_***T***_3_*.

### 2.3 RMSD Calculation

For each generated pose **T**_*final*_, its quality is assessed by calculating the Root Mean Square Deviation (RMSD) of the ligand’s C*α* atoms relative to a known ground-truth reference structure. Let 𝒞_*l*_ = {**c**_1_, …, **c**_*m*_ }be the initial coordinates of the ligand’s C*α* atoms and 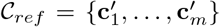 be the coordinates in the reference structure. The RMSD is calculated as:

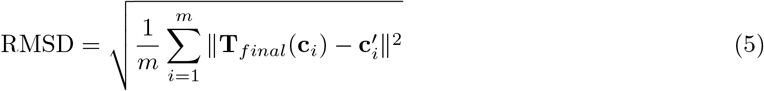

## 3 Results

To validate our docking pipeline, we performed a docking experiment on a well-characterized protein complex, 1AVW, consisting of a receptor (chain A) and a ligand (chain B). The performance of the algorithm was evaluated by calculating the ligand C*α*-RMSD against the native structure.

### 3.1 Experimental Setup

The docking simulation was executed using the final parameters determined through iterative testing. Specifically, the number of concave or convex features (‘`num features` ‘) was set to 50, the MSMS surface density (‘msms density’) was set to 0.5, and the local patch radius (‘`patch radius` ‘) was set to 6.0 Å. These parameters were chosen to balance computational efficiency with a comprehensive search of the conformational space.

The total number of initial docking trials was 5000 (50 × 50 concave-convex pairs and 50 × 50 convex-concave pairs). The entire simulation, from surface generation to the completion of all 5000 alignment optimizations, was completed in approximately 2 minutes and 40 seconds on a standard Apple ARM64 processor, demonstrating the high computational efficiency of the approach.

### 3.2 Docking Performance and Analysis

Our method successfully identified a high-quality near-native docking pose. The best-ranked pose, as determined by RMSD to the native ligand position, achieved a value of **3.43 Å**. This result is considered a “medium” to “good” quality prediction according to the CAPRI (Critical Assessment of PRedicted Interactions) criteria, validating the efficacy of our geometric docking algorithm.

Table 1 presents the top 10 docking poses sorted by their C*α*-RMSD. It is noteworthy that the geometric alignment score, while effective for local patch optimization, does not serve as a reliable global scoring function for discriminating near-native poses. For instance, the pose with the lowest RMSD (3.43 Å) had a geometric score of 1.818, whereas the pose with the highest geometric score (2.676) had a very poor RMSD of 25.71 Å. This highlights a key challenge in protein docking: the need for a more physically realistic scoring function to complement the geometric search.

**Table 1.**
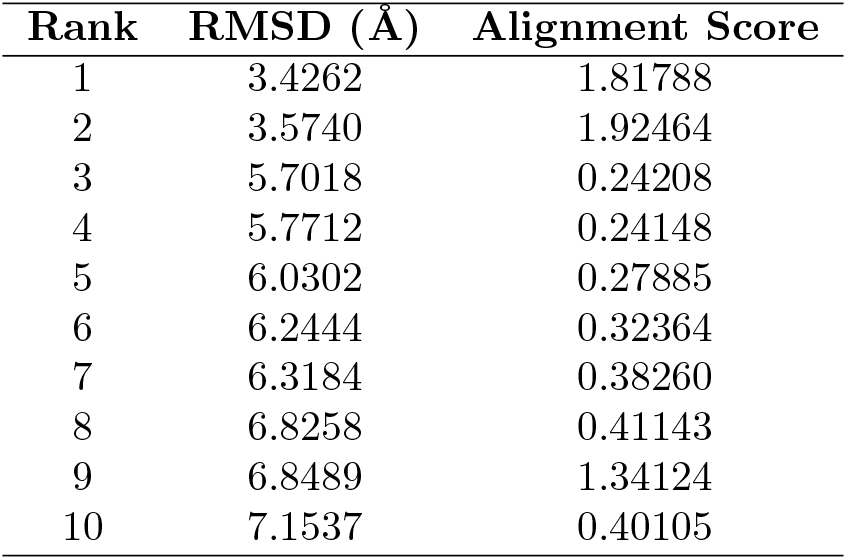
Top 10 Docking Results for 1AVW Sorted by RMSD.

| Rank | RMSD (Å) | Alignment Score |
| --- | --- | --- |
| 1 | 3.4262 | 1.81788 |
| 2 | 3.5740 | 1.92464 |
| 3 | 5.7018 | 0.24208 |
| 4 | 5.7712 | 0.24148 |
| 5 | 6.0302 | 0.27885 |
| 6 | 6.2444 | 0.32364 |
| 7 | 6.3184 | 0.38260 |
| 8 | 6.8258 | 0.41143 |
| 9 | 6.8489 | 1.34124 |
| 10 | 7.1537 | 0.40105 |

### 3.3 Visual Confirmation

Visual inspection confirms the quality of the top-ranked pose. Figure 1 shows a superposition of the lowest-RMSD docked ligand (3.43 Å) onto the native crystallographic structure. The model accurately recapitulates the binding mode and orientation of the ligand in the receptor’s binding site.

**Figure 1.**
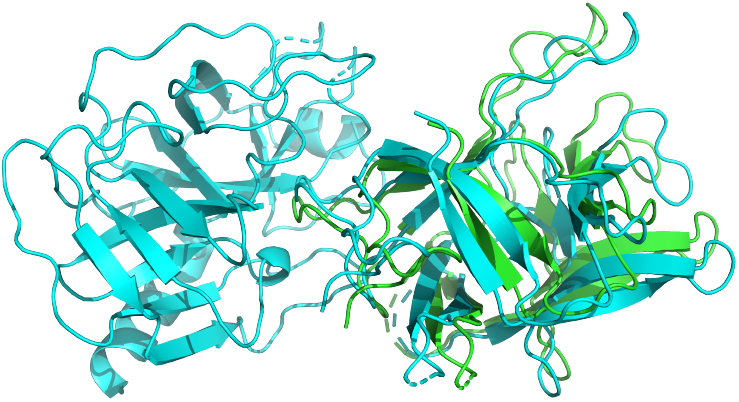
Superposition of the best-ranked docked pose (in blue) onto the native crystallographic ligand (in green). The receptor is shown as a grey surface. The model achieves an RMSD of 3.43 Å.

## 4 Discussion

The primary goal of this study was to develop and validate an efficient algorithm for protein-protein docking by leveraging modern mathematical frameworks. The successful identification of a 3.43 Å RMSD pose for the 1AVW complex demonstrates the fundamental viability of our approach. The key innovation lies in the synergy between a robust geometric feature detection method and a differentiable local alignment module that operates directly on the Lie group SE(3) guided by an Optimal Transport score.

The efficiency of the pipeline is a significant advantage. Completing a 5000-trial docking simulation in under three minutes suggests that the method is well-suited for large-scale virtual screening applications. This speed is achieved by the efficiency of the k-d tree neighbor search and the rapid convergence of the gradient-based SE(3) twist optimization, which is a direct benefit of the smooth, unconstrained optimization landscape provided by the Lie algebra representation.

### 4.1 Limitations and Future Directions

Despite its success in rapidly generating near-native poses, LieOTDock has several limitations that open avenues for future research. The most significant shortcoming is the scoring function. As shown in Table 1, the geometric alignment score used for local refinement does not strongly correlate with the global RMSD. This indicates that while our method is an excellent pose generator, the scoring function is not yet sufficient to reliably identify the most near-native pose from the generated candidates. The current score, based on shape complementarity and a simple clash penalty, lacks the physical realism required for accurate final-stage ranking. The field of protein docking has seen a shift towards more sophisticated, physics-based and machine learning-based scoring functions that incorporate terms for van der Waals interactions, electrostatics, and solvation energy [4]. Integrating such a scoring function, perhaps one based on graph neural networks like GNINA [6], as a post-processing or re-ranking step is a critical priority for future work.

Another limitation is the assumption of rigid-body docking. Proteins are inherently flexible, and conformational changes upon binding are common. Our current framework does not account for this flexibility. An exciting future direction would be to explore the use of Unbalanced Optimal Transport (UOT) [11]. UOT is a generalization of OT that allows for the comparison of distributions with unequal mass. In the context of docking, this could potentially model flexible backbones or side chains where parts of a surface patch might move, or be inserted or deleted, during binding. The challenge would lie in formulating an appropriate mass penalty term within the UOT framework and managing the increased computational complexity.

Finally, the initial feature sampling, while robust, could be improved. The current method relies on local geometry but does not incorporate any chemical or evolutionary information. Integrating information about residue conservation, hydrophobicity, or electrostatic potential into the feature scoring could help to prioritize biologically relevant interfaces and further improve sampling efficiency.

## 5 Conclusion

We have developed and successfully implemented LieOTDock, a novel and highly efficient pipeline for protein-protein docking that is grounded in the mathematics of Lie groups and Optimal Transport. Our method introduces a robust geometric feature scoring algorithm that overcomes the limitations of discrete surface descriptors. By combining this with a differentiable local alignment strategy on SE(3), our algorithm successfully generated a near-native pose of 3.43 Å for the 1AVW complex in a remarkably short time. The primary strength of the current method is its speed and its effectiveness as a pose generator. While the current scoring function is a limitation, it also means that LieOTDock provides a powerful geometric sampling engine that can be readily combined with more advanced, physics-based or AI-driven scoring functions. Future work will focus on this integration, as well as exploring extensions to handle protein flexibility, to further enhance the power and applicability of the LieOTDock framework.

## AI Declaration

The authors acknowledge the use of Gemini, a large-scale language model from Google, for assistance in literature search, code implementation, debugging, and manuscript polishing. The overall research plan, core algorithmic design, and final conclusions were directed by the human authors, who take full responsibility for the content of this work.

## Code Availability

The source code for LieOTDock is openly available on GitHub at https://github.com/YueHuLab/LieOTDock.

